# Evaluation of orange pulping residues as an alternative growth medium for *Thraustochytrium* sp

**DOI:** 10.64898/2026.05.25.727442

**Authors:** Joaquín Ramos Céspedes, Sergio Castillo Fernández-Dávila, Rocío Navarro Segura, Yazmín Dumet Poma, Siomi Muñoz Titto

## Abstract

In this study, the lipid content produced by *Thraustochytrium* sp. in a medium prepared from orange pulp residue was compared with that obtained in a conventional medium. The pulp residue was subjected to freezing, blending, and filtration in seawater to prepare three treatments: conventional medium (T1), filtrate (T2), and filtrate supplemented with KNO_3_ (T3). A growth curve was performed over six days, after which biomass and lipid content were analyzed. The results showed that T2 exhibited the highest cell growth and biomass yield (5.24 g/L). However, lipid content was higher in the conventional medium (38.75%), whereas the treatments containing orange extract showed lower values. These findings suggest that the medium prepared from orange pulp residue is feasible for the growth of *Thraustochytrium* sp., but requires optimization to enhance lipid accumulation and its potential use in sustainable bioprocesses.

## INTRODUCTION

The global demand for high value-added lipids has increased significantly, as they are becoming part of consumers’ daily diets. Among these, long-chain omega-3 polyunsaturated fatty acids (PUFAs), such as eicosapentaenoic acid (EPA) and docosahexaenoic acid (DHA), are of particular interest. According to Valenzuela *et al*. (2011), these fatty acids are found in high amounts in fish and fish oil, which are widely used as nutritional supplements due to their proven effectiveness in the prevention and treatment of cardiovascular, neurodegenerative, cancer-related, and immune-associated diseases.

In this context, microalgae such as *Thraustochytrium* sp. have been identified as a promising alternative source of fatty acids. Given current challenges such as overfishing and global warming, they represent a more sustainable option (Barta *et al*., 2021). However, the high cost of conventional culture media remains a major limitation for industrial-scale production.

In Peru, large quantities of orange peel waste are generated and commonly discarded as household waste, creating an environmental concern (Negro *et al*., 2017). Nevertheless, citrus peels are rich in carbon and nitrogen sources and have been shown to enhance microalgal growth and composition (Nateghpour *et al*., 2021). Therefore, this study aimed to evaluate a novel culture medium incorporating orange processing residues to improve lipid yield in *Thraustochytrium* sp. cultivation.

## MATERIAL AND METHODS

### Microalgal strain

The *Thraustochytrium* sp. strain was obtained from the strain bank of the company Biorefinerías del Perú S.A.C. It was maintained in the company’s conventional medium under constant orbital agitation at 150 rpm and at a temperature of 21 °C. These cultures were used as inoculum for the different experimental treatments.

### Production of orange extract

Orange residues were initially stored in a freezer. Subsequently, they were thawed and 200 g of the residue were weighed and blended with 1 liter of filtered seawater. The resulting slurry was left to stand for 24 hours, after which the solid fraction was separated by filtration using a strainer.

### Preparation of treatments

For the evaluation of the filtrate, three treatments were prepared. The first treatment (T1) corresponded to the control medium, whose formulation is confidential to Biorrefinerías del Perú S.A.C. For the other two treatments, the filtrate was used directly (T2), and in treatment T3 it was supplemented with potassium nitrate (6 g/L) as a nitrogen source. The pH of these two treatments was adjusted to the pH of the conventional medium (pH = 6). The media were prepared in 250 mL flasks filled to 80% of their capacity and sterilized by autoclaving. Finally, the media were inoculated with 10% (v/v) of the mother cultures.

### Analysis of treatments

Growth performance was evaluated by generating growth curves based on daily cell counts using a Neubauer chamber for at least five days of cultivation. Biomass production was assessed after approximately five days by harvesting cultures through centrifugation at 4000 rpm for 5 minutes, followed by washing and drying of the recovered biomass to determine dry weight.

Lipid extraction and quantification were performed using a method similar to that proposed by Bligh and Dyer (1959). Dry biomass was ground, and 0.1 g was mixed with 0.1 g of aluminum oxide and 2 mL of a chloroform:methanol (2:1) solution. The mixture was homogenized by vortexing for 5 minutes and centrifuged at 4000 rpm for 5 minutes, recovering the supernatant. This step was repeated with an additional 2 mL of solvent until a clear supernatant was obtained. Subsequently, 3 mL of 0.5% MgCl_2_ solution were added, vortexed for 5 minutes, and centrifuged at 4000 rpm for 5 minutes. The organic phase was recovered using a Pasteur pipette and dried at 60 °C until solvent evaporation. Lipid content was calculated by dividing the lipid mass obtained by 0.1 g of dry biomass.

### RESULTS AND DISCUSSION

The accumulation of lipids in oil-producing microorganisms is strongly influenced by the composition of the culture medium and by the nature of the available carbon and nitrogen sources, factors that regulate both cell growth and metabolic pathways towards lipogenesis. In thraustochytrids such as *Thraustochytrium* sp., lipid biosynthesis is closely associated with nutrient availability and carbon/nitrogen balance, factors that determine the shift between cellular proliferation and lipid accumulation (Costa *et al*., 2024). Recent studies also highlight that the use of alternative substrates, including agroindustrial waste as carbon sources, can modify the metabolism and lipid production efficiency of oleaginous microorganisms, suggesting the need to optimize these sources for sustainable biotechnological processes (Ringel *et al*., 2025).

Furthermore, studies with thraustochytrids have shown that the composition of the culture medium has significant effects on the organism’s physiology and lipid accumulation, supporting the idea that variations in carbon sources can trigger contrasting metabolic responses in the microalgae studied (Paredes *et al*., 2025). In this context, the results obtained in the present study are presented and discussed to analyze the trends in growth, biomass production and lipogenesis under the evaluated experimental conditions.

In the comparison of the three treatments, it was observed that treatment T1 (control medium) reached the stationary phase on day four, whereas T2 and T3 (both containing orange peel extract) were still in the exponential phase. This suggests that in T1 the available nutrients were depleted more rapidly, which may have accelerated the transition to the stationary phase and triggered a metabolic shift from cellular growth toward lipid accumulation. This behavior has been widely described in thraustochytrids, where nitrogen limitation acts as a key signal inducing lipogenesis (Xiao *et al*., 2018).

In contrast, in T2 and T3, the presence of orange peel extract likely provided a more sustained nutrient availability, prolonging the exponential phase and primarily promoting cellular proliferation. This phenomenon is consistent with the findings reported by Jung *et al*. (2020) in studies with *Schizochytrium* sp. ABC101, where lipid accumulation was shown to be tightly regulated by nitrogen balance; conditions with higher nutrient availability promote cellular growth, whereas nitrogen limitation redirects metabolism toward lipid storage.

Microscopic observations revealed significant morphological differences between treatments. In T1, the cells were larger and showed intracellular accumulations consistent with lipids, while in T2, the cells were smaller and tended to form aggregates or clusters, suggesting a possible cellular stress response or partial growth inhibition (Chen *et al*. 2022).

These observations correlate with the yield differences in Figure 2. The lower yield observed in T1 can be explained by the fact that this treatment reached the stationary phase more quickly due to early nutrient depletion. Although glycerol, an input that is part of the composition of the control treatment, is a pure and easily assimilable carbon source, its rapid availability would have limited the effective time for biomass accumulation, favoring mainly the accumulation of intracellular lipids rather than cell proliferation. The high biomass yield value in T2 and T3, compared to T1, suggests a more sustained availability of nutrients, probably associated with the complex composition of the orange peel extract. This condition would have prolonged the cell growth phase, resulting in higher total biomass production, even though carbon was not supplied in a totally efficient manner. However, these differences in biomass were relatively small.

**Figure 1.**
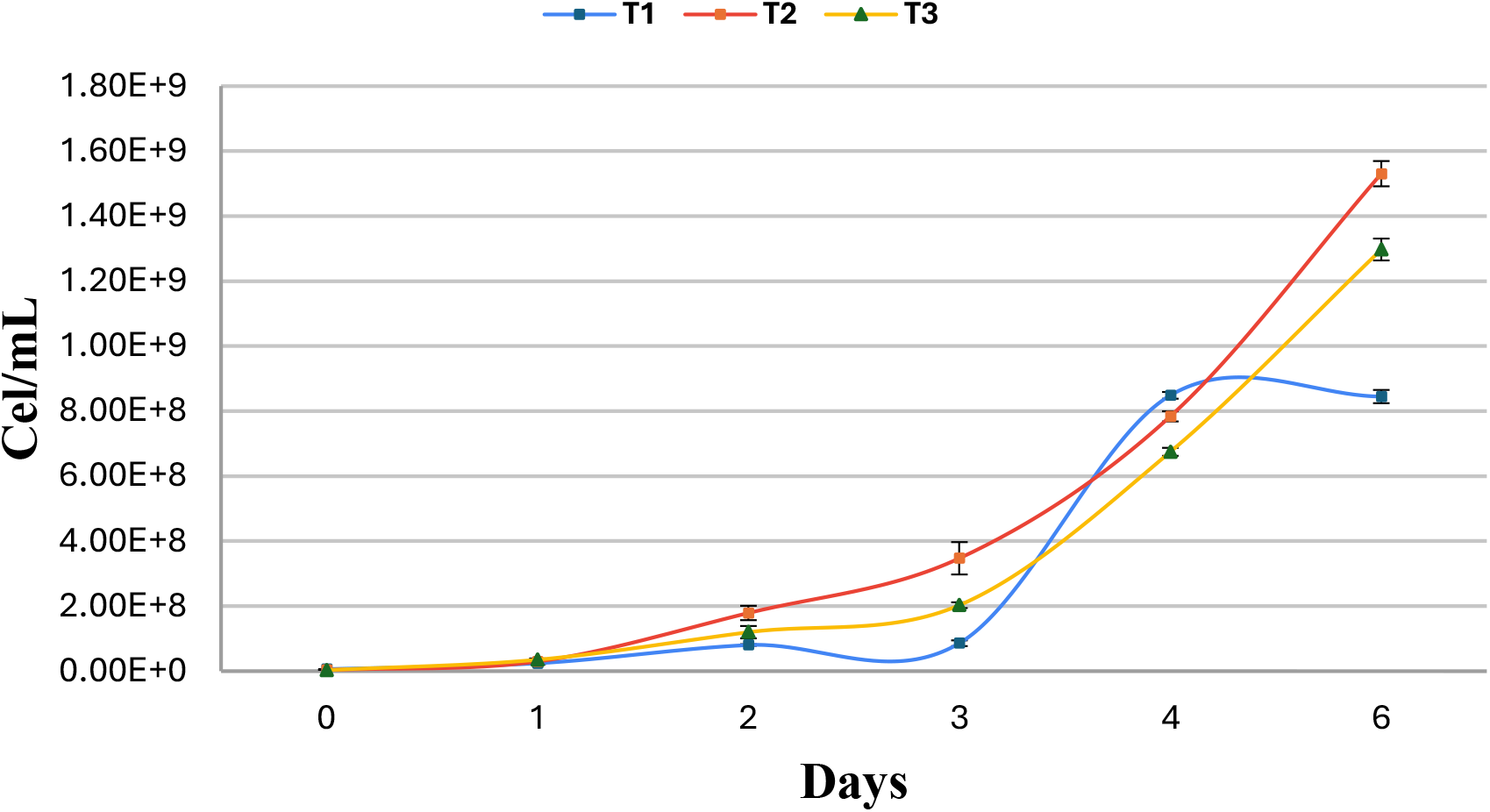
Growth curves based on cell counting of *Thraustochytrium* sp. under different culture treatments *Note*. T1: Conventional medium; T2: filtrate of orange peel extract; T3: filtrate supplemented with KNO_3_.

**Figure 2.**
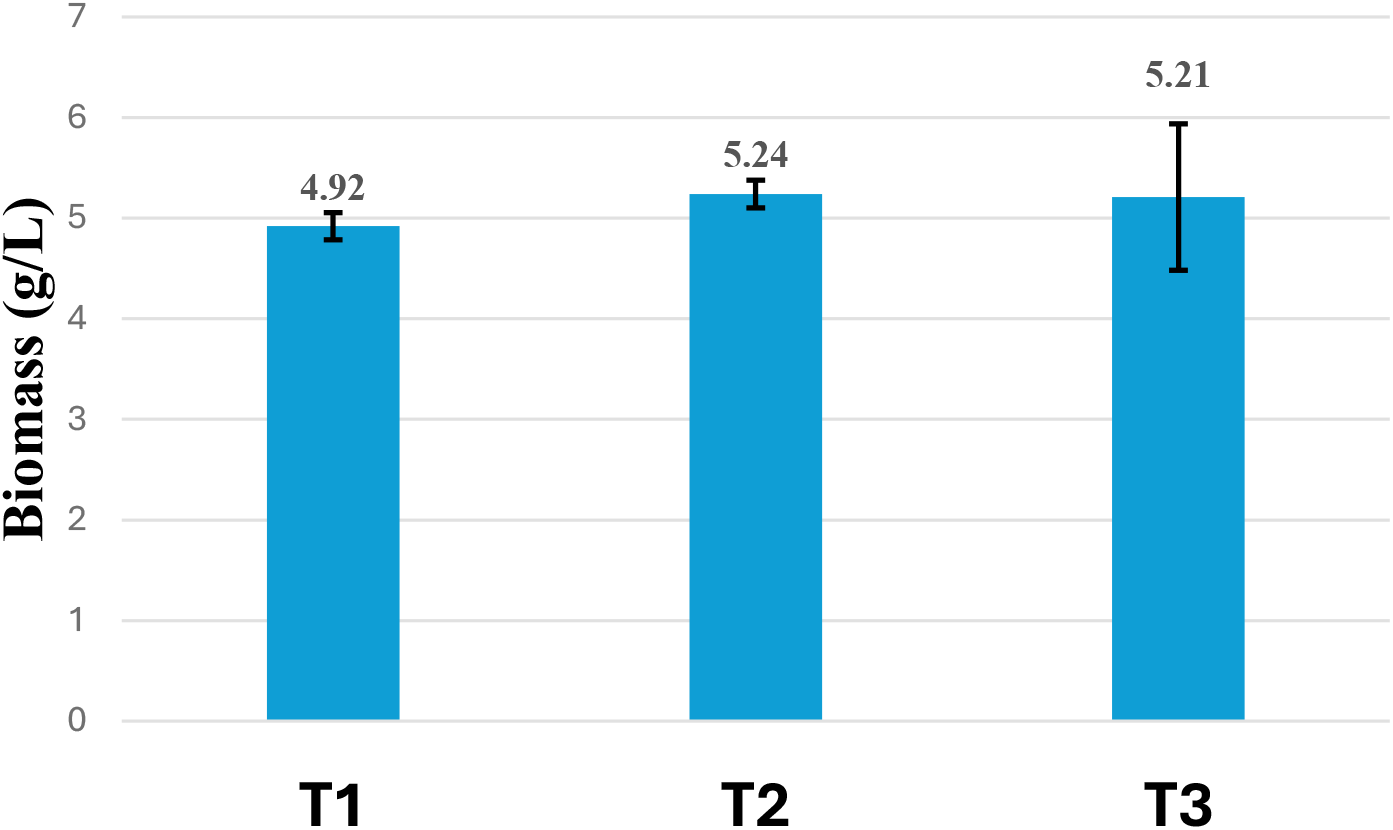
Biomass production in the treatments *Note*. T1: Conventional médium; T2: filtrate of orange peel extract; T3: filtrate supplemented with KNO_3_.

Regarding lipid extraction yield relative to biomass (figure 3), Treatment 1 showed a yield of 38.7%, whereas Treatments 2 and 3, formulated with orange extract, exhibited significantly lower values of 8.25% and 6.67%, respectively. These results reinforce the idea that lipid production is being affected by the nutrients present in the culture medium, specifically those provided by the orange extract. In addition to factors such as temperature, pH, salinity, and others, nutrient composition is one of the main determinants influencing fatty acid production. Thraustochytrids are known to increase lipid synthesis when exposed to stress conditions, such as temperatures outside the optimal range or limited nitrogen availability. In particular, a high carbon-to-nitrogen (C/N) ratio significantly enhances the production of docosahexaenoic acid (DHA) (Patel *et al*., 2021).

**Figure 3.**
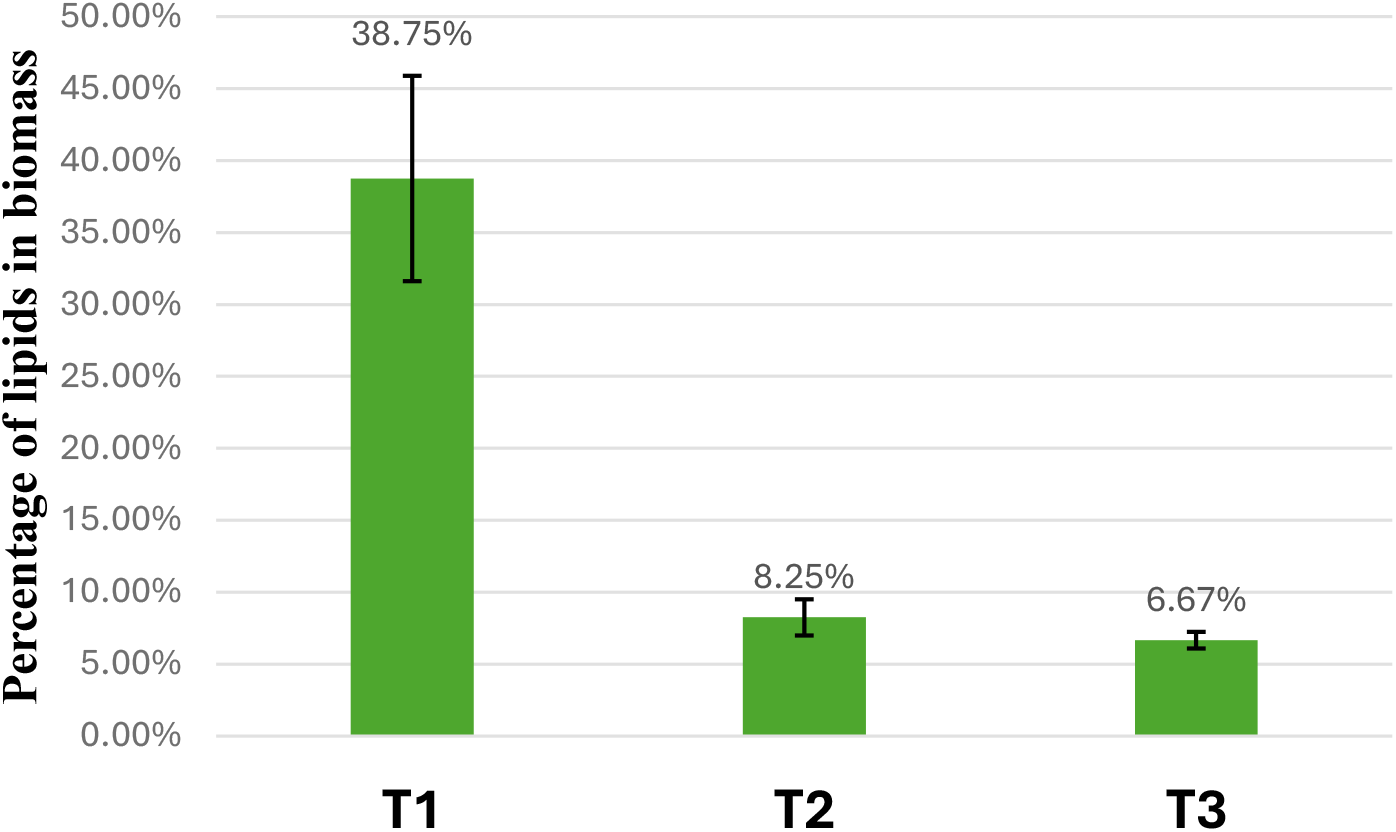
Lipid Content in Biomass (%) *Note*. T1: Conventional médium; T2: filtrate of orange peel extract; T3: filtrate supplemented with KNO_3_.

In other studies evaluating the influence of culture conditions—such as carbon and nitrogen sources—on species like *Thraustochytrium striatum*, it was reported that a low C/N ratio favored cellular growth, whereas a high C/N ratio promoted fatty acid accumulation. A maximum lipid content of 25% (g/g DCW) was obtained with a glucose/YEP ratio of 30:1 (w/w) (Xiao *et al*., 2018).

Considering the importance of medium optimization for lipid production, it is likely that the lack of precise information regarding the carbon and nitrogen concentrations in the orange extract—and therefore the C/N ratio—resulted in insufficient control of this critical parameter, which is essential for efficient lipid production.

## CONCLUSION

The culture medium formulated from orange filtrate supported the cellular growth of *Thraustochytrium* sp. Among the treatments evaluated, the highest biomass production was observed in treatment T2 (orange filtrate alone), followed by T3 (orange filtrate supplemented with potassium nitrate), and finally the control treatment (T1). Although T2 achieved the greatest biomass yield (5.24 g/L), it exhibited a relatively low lipid proportion, which may be attributed to an imbalance in the carbon-to-nitrogen (C/N) ratio. In contrast, T1, despite showing the lowest cell growth, recorded the highest average lipid content (38.75%). Overall, these findings indicate that orange extract represents a promising and viable substrate for enhancing cellular growth. However, if the primary objective is to maximize lipid production, careful optimization of nutrient availability is required.

## Acknowledgments

To the Office of the Vice-Rector for Research of UNALM for the funding granted through the XIII Call for Research Project Funding in Research Circles 2024; to our advisor, Patricia Angélica Moreno Díaz and to Biorefinerías del Perú S.A.C. for providing us with access to their facilities and the necessary equipment.

